# Competing Effects of Agonist and Antagonist Vibration on the Proprioceptive Sense of Force

**DOI:** 10.64898/2026.02.05.704112

**Authors:** Kaitlyn G Sutton, Olivia R Ryan, Gregory E P Pearcey

## Abstract

Motor unit (MU) firing is affected by motoneuronal persistent inward currents (PICs), which heavily contribute to gain control of motor output. PICs are highly sensitive to inhibition; for instance, Ia reciprocal inhibition via antagonist muscle vibration drastically reduces discharge rate hysteresis (ΔF), an estimate of PIC magnitude. A direct link between sensitivity of PICs to inhibition and voluntary force control, however, has not been established. To determine whether force control is altered with inhibition of PICs, we recorded high-density surface EMG from the tibialis anterior, while 11 participants (5F; 6M) completed and isometric force reproduction task. Tendon vibration was applied to the agonist or antagonist muscle during the first (with visual feedback) or second contraction (without visual feedback) and participants were asked to match percieved effort across contractions, in an attempt to match neural drive to the motor pool. In support of our hypothesis, torque and MU firing rates were reduced when vibration was applied to the antagonist (torque: *p* < .0001; MU firing rate: *p* < .0001), but not agonist (torque: *p* = .9980; MU firing rate: *p* = .312) muscle tendon in the second contraction, compared to control. Conversely, when vibration was applied during the first contraction, opposite effects were observed. These results suggest that PICs play a role in the proprioceptive sense of force, offering a potential link between PICs and voluntary force control, which may be important for understanding and treatment of motor impairments.

**KEY POINTS:**

- Motoneuronal persistent inward currents amplify synaptic currents and therefore heavily contribute to motor output, however they are extremely sensitive to Ia reciprocal inhibition induced by muscle tendon vibration.
- We show that modulation of PICs severely impacts human force sense using an effort-based force reproduction paradigm which enabled us to manipulate combinations of tendon vibration and visual feedback.
- These findings provide a link between PICs and functional motor output, which may be important for understanding neurological impairments and informing rehabilitation strategies.

## INTRODUCTION

All movements are generated by the controlled activation of motor units (MUs), which consist of motoneurons and the muscle fibers in which they innervate (Sherrington, 1906). Motoneurons are unique because they can be studied non-invasively in humans due to their one-to-one spike ratio with the fibers they innervate, meaning that firing patterns of motoneurons and their MU are the same (Heckman and Enoka 2012). Motoneurons integrate combinations of excitatory, inhibitory and neuromodulatory inputs (i.e. serotonin; 5-HT and norepinephrine; NE), which have powerful effects on the intrinsic excitability of motoneurons through the facilitation of persistent inward currents (PICs; Heckman et al., 2009). Activation of PICs can amplify synaptic currents up to 3-5-fold (Lee & Heckman, 1996, 2000), but their facilitation via monoaminergic pathways is diffuse and simultaneously affects multiple motor pools within a limb (Skagerberg & Björklund, 1985). Interestingly, the activation of PICs also increase sensitivity to inhibition, particularly Ia input from the antagonist muscle (Hyngstrom et al., 2007), whereas Ia monosynaptic input to agonists has little impact on PIC amplitude (Heckman & Lee, 1999).

High-density surface electromyography (HD-sEMG) combined with blind source separation decomposition algorithms has enhanced the ability to sample tens of simultaneously active MUs (Holobar et al., 2009; Negro et al., 2016; Del Vecchio et al., 2020), which has helped glean incredible insights about motor output in various conditions. Similar to agonist (Lapole et al., 2023), when antagonist tendon vibration is applied, estimates of PIC magnitudes (using the paired motor unit analysis technique; Gorassini et al., 2002) are reduced during both plantar- and dorsiflexion contractions (Pearcey et al., 2022). Interestingly, during the antagonist vibration experiment, participants anecdotally reported that they had more difficulty reaching the target force when antagonist vibration applied compared to conditions without vibration. This may have been due to a requirement for enhanced synaptic input to offset the vibration-induced inhibition, which may have also minimized the effect of vibration on PIC estimates. To better understand the relationship between PICs and motor function, we can link this altered excitability to humans’ force sense.

The sense of force is a component of proprioception and is the ability to correctly perceive a given level of force (Proske & Gandevia, 2012). It can be measured using a force reproduction task, which requires reproducing a force with the same limb in the absence of performance feedback (Proske and Allen, 2019). This method is said to rely primarily on proprioceptive inputs from muscle spindles and Golgi tendon organs (Monjo et al., 2018), as well as sensory inputs from mechanoreceptors in the skin (De Serres and Fang 2004; Proske and Allen 2019), which informs the required force in the absence of feedback.

Given PICs sensitivity to reciprocal Ia inhibition and the increased perceived difficulty with applied antagonist tendon vibration, the purpose of this study was to explore the influence of the addition as well as the removal of either agonist or antagonist vibration on the force output and motor unit firing behaviour of the agonist muscle, in conditions where neural drive remains constant. To achieve conditions of similar neural drive, we used the force reproduction task, by asking participants to focus on replicating their effort in the task with visual feedback, after feedback was removed. Due to the Ia afferent discharge and subsequent reciprocal inhibition elicited by antagonist vibration applied to the antagonist muscle tendon, we hypothesized that force production would require greater neural drive to reach the same target force with concurrent vibration. Therefore, in a force reproduction task, force output would be reduced during antagonist vibration and increased following removal of antagonist vibration when participants attempted to reproduce their perceived effort without visual feedback of their force output. Given that agonist Ia monosynaptic input has little impact on PIC amplitude (Heckman & Lee, 1999), we hypothesized minimal effects on force reproduction in the presence and following removal of agonist vibration.

## METHODS

### Participants and Ethical Approval

Both females (n = 5; 23.2 ± 3.6 yr.) and males (n = 6; 22.2 ± 2.4 yr.) were recruited to participate in this study. Participants were apparently healthy, such that they have no history of neuromuscular, cardiovascular, or metabolic impairment and do not take medications which affect their motor abilities. All participants provided written informed consent to the experimental protocol prior to partaking in any lab procedures, which was approved by the Interdisciplinary Committee on Ethics in Human Research at Memorial University (ICEHR #: 20240936-HK).

### Experimental Apparatus

#### Torque

Participants were seated comfortably in a HUMAC NORM isokinetic dynamometer (CSMi, Stoughton, MA) and their right foot was secured in a footplate attachment with the ankle at an angle of 95° and the knee at an angle of 20° of flexion (see fig 1). Velcro straps were wrapped around the foot to secure it to the plate and prevent extraneous movement. Shoulder straps secured participants to the chair to minimize changes in body position. A target line along with visual torque feedback was provided to participants during respective contractions on a 35-inch monitor placed ∼1.5 m in front of the participant at eye level using custom written MATLAB software. Torque signals were sampled at 2,048 Hz and smoothed offline with a 10-Hz low-pass filter (fifth-order Butterworth filter).

**Figure 1.**
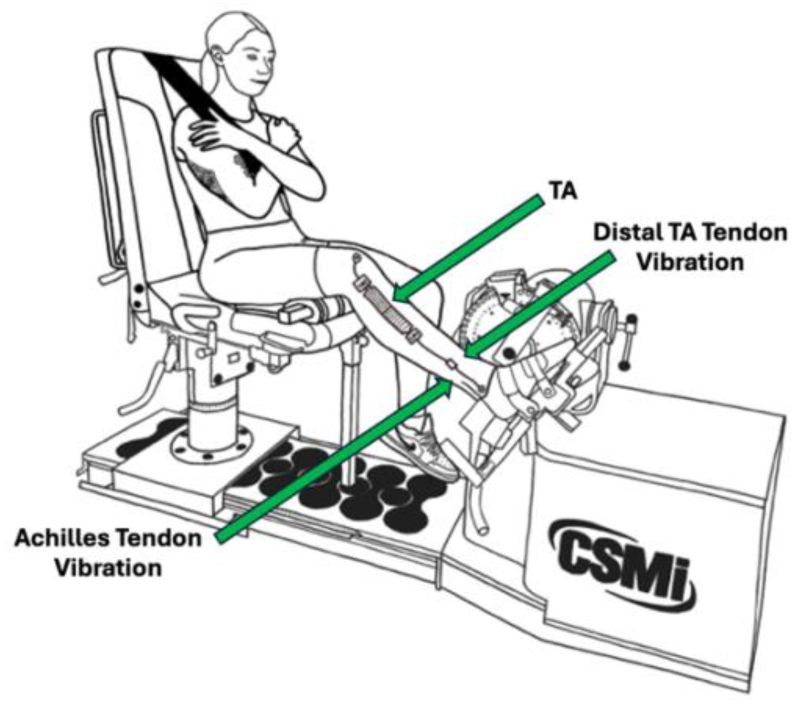
Experimental setup.

#### High-density surface electromyography (HD-sEMG)

Prior to electrode placement, the skin was carefully prepared by removing excess hair and gently rubbing the area over the target muscle with an abrasive and conductive gel (NUPREP, D.O. Weaver and Co., Aurora, CO, USA). HDsEMG signals were collected from the TA (right leg), a dorsiflexor muscle of the ankle, using two electrode arrays (M8X4D10; 8 x 4 configuration with 10mm I.E.D.; ReC Bioengineering, Turin, Italy) fixed to the skin over the muscle belly. Hypafix tape (BSN Medical Inc., Hamburg, DE) was used to mitigate movement of the arrays. We securely fixed two Ag/AgCl electrodes (Cardinal Health, Dublin, OH, USA) to the patella and the lateral malleolus of the right leg as reference electrodes. HD-sEMG signals were amplified (192x) and acquired (2048Hz) using MEACS amplifiers (LISiN, ReC Bioengineering, Politecnico di Torino, Turin, Italy).

#### Vibration

A commercially available handheld massager (Thrive Handy Massager, THRIVE, Japan) was used to apply vibration at 108 Hz to either the tibialis anterior tendon (agonist) or the Achilles tendon (antagonist) of the contracting muscle. The massager was manually positioned and held perpendicularly to the targeted tendon and vibration commenced 2 s before the ascent of respective contractions and ceased 2 s after the descent.

### Experimental Protocol

#### Maximum voluntary contractions

Once setup was complete, real-time HD-sEMG signals were examined to ensure high quality recordings (i.e. a high signal-to-noise ratio). Participants performed a minimum of two maximum voluntary contractions (MVCs) of their dorsiflexors lasting four seconds each, with two minutes of rest between each contraction to mitigate effects of fatigue. Members of our lab team gave verbal encouragement during MVC collection to ensure maximal effort. If the two peak torque values generated during the MVCs significantly differed (i.e. > 5% apart), a third MVC was performed. If the two peak torque values generated were within 5% of each other, the mean peak torque value was used for subsequent xnormalization of all trials.

#### Experimental conditions

Once the target of dorsiflexion force was determined, participants completed a series of force reproduction trials consisting of two contractions each. All contractions followed a trapezoidal shaped ramp with a three second ascent, six second hold and three second descent and reached 25% of participants’ MVC.

Participants completed two sets of five trials each with different vibration conditions for a total of ten force reproduction trials (20 total contractions). The control condition consisted of dorsiflexion with no vibration. Antagonist vibration conditions involved dorsiflexion with vibration applied to the Achilles tendon, either during the first contraction only, or the second contraction only. Agonist vibration conditions involved dorsiflexion with vibration applied to the distal TA, either during the first contraction only, or the second contraction only (see fig. 2). Before beginning the experimental conditions, participants were required to practice each condition until they were both comfortable and proficient. Participants were given one minute to rest between trials.

**Figure 2.**
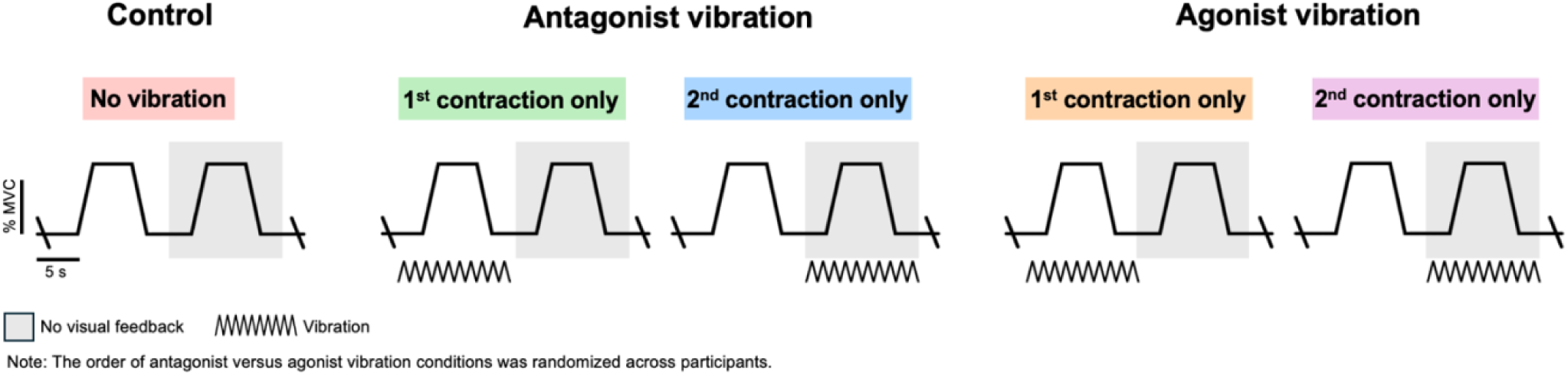
The five experimental conditions.

#### Visual feedback and perceived effort

Regardless of the experimental condition, participants were given visual feedback during the first contraction within each trial but were not given visual feedback during the second contraction of each trial (see above; fig 2). Participants were instructed, with emphasis, to use the exact same effort during both contractions within a given trial, regardless of whether one felt easier or more challenging than another. These instructions were reiterated to participants throughout the study to ensure consistent effort. Further, participants were asked to rate their perceived level of effort required to perform the task during the hold of each contraction. A visual analog scale from 0 to 10 was used, where 0 meant no effort and 10 meant absolute maximal effort (see fig 3).

**Figure 3.**
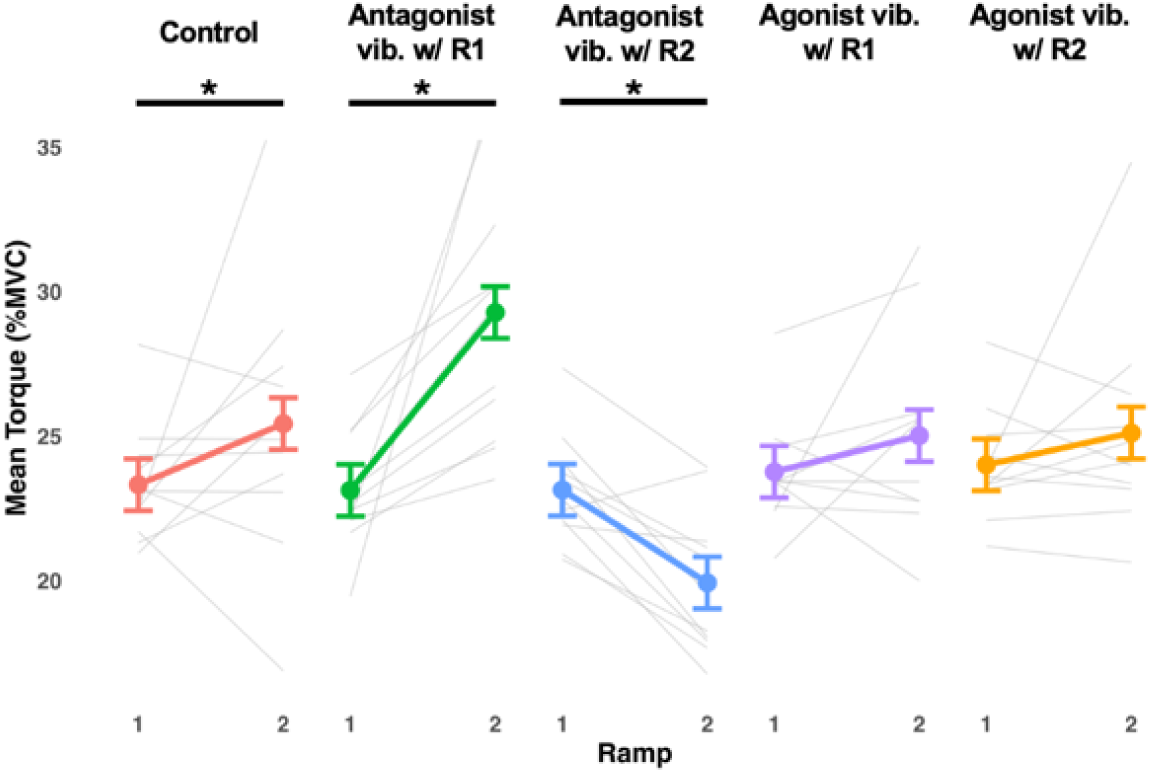
Mean torque (% MVC) values for the tibialis anterior (TA) during each contraction within each experimental condition. Points represent the model estimates from the linear mixed effects model and the error bars are the corresponding 95% confidence intervals. Grey lines in the background of each condition indicate individual participant behaviour. Asterisks indicate significant difference in mean torque between contractions (p < .05).

### Data Analysis

#### Torque data

Following data collection, torque data was filtered with a low pass 20Hz filter to remove any high frequency artifacts. Any signals with significant noise were omitted from the results. The mean torque of each contraction was determined from the middle 3 seconds of the instructed plateau.

#### Error calculations

To further assess force reproduction accuracy, we quantified absolute error (AE) as the absolute value of the difference between the reproduced torque and the target torque. Additionally, to investigate directional bias, constant error (CE) was calculated as the signed difference between the reproduced torque and the target torque, where positive values indicate an overshoot and negative values indicate an undershoot. Both AE and CE were expressed as a percentage of the target torque to normalize error across participants.

#### Motor unit decomposition

HD-sEMG signals were first bandpass filtered (10-500Hz) and then manually inspected. Channels with considerable noise or artifacts were omitted from further analysis. The remaining HD-sEMG signals were then decomposed using the Convolution Kernel Compensation (CKC) algorithm (Holobar & Zazula, 2007; Holobar et al., 2009), which uses convolutive blind-source separation methods (Negro et al., 2016) within MATLAB (MATLAB (R2023b), The MathWorks Inc., Natick, MA). After decomposition, individual motor unit spike trains were inspected for accuracy and edited using the CKC inspector within DEMUSE to correct any errors made by the decomposition algorithm. Instantaneous firing rates were quantified as the inverse of the interspike interval. To generate smooth estimates of firing rates, we used a support vector regression model (Beauchamp et al., 2022) with custom written MATLAB scripts. Similar to the torque, the mean firing rate of motor units was taken from the middle 3s of each instructed plateau of each contraction.

### Statistical Design

Statistical analysis was performed using R statistical software (v4.1.2; R Core Team 2021). For each dependent variable of interest (i.e. torque, AE, CE and MU firing rate) linear mixed-effects models were fit using the lme4 package (Bates et al., 2015). Condition and contraction were included as fixed effects, with a random intercept for participant (PID). For MU firing outcomes, a nested random effect for MUID within participant was added to account for repeated measures. Model fit was examined using likelihood ratio tests (LRTs), such that all main models were compared against reduced models with condition, contraction, or their interaction terms removed. These comparisons are reported as χ^2^ statistics with associated degrees of freedom and p-values.

For all mixed models, we confirmed that that assumptions of linearity, homoscedasticity and normality of residuals were valid for all data. For main effects or interactions found to be significant, estimated marginal means (EMMs) and pairwise contrasts were generated using the emmeans package (Lenth, 2022). For these post-hoc tests, Satterthwaite’s method was used to estimate degrees of freedom and significance using the ImerTest package (Kuznetsova et al., 2017), and Tukey adjustment was applied for multiple comparisons. Effect sizes (Cohen’s d with 95% CI) were calculated from model residual variance for all pairwise contrasts. All data were visualized using ggplot2 (Wickham, 2016).

## RESULTS

### Motor unit identification

A total of 589 MU spike trains were identified and tracked across contraction pairs. We identified 121 MUs with a mean of 11 ± 4.88 MUs per participant per trial in the control condition; 114 MUs (10.36 ± 4.06 MUs per participant per trial) with agonist vibration applied in the first of two contractions; 116 MUs (10.55 ± 3.11 MUs per participant per trial) with agonist vibration applied in the second of two contractions; 121 MUs (11 ± 4.12 MUs per participant per trial) with antagonist vibration applied in the first of two contractions; and 117 MUs (10.64 ± 4.23 MUs per participant per trial) with antagonist vibration applied in the second of two contractions.

### Torque

Condition (χ^2^(8) = 61.78, *p* < .0001), contraction (χ^2^(5) = 42.44, *p* < .0001) and their interaction (χ^2^(4) = 35.35, *p* < .0001) were significant predictors of torque.

#### Differences between conditions within each contraction (compared to control)

During the first contraction in each pair, torque did not significantly differ between control and any of the vibration conditions. This was completely expected since they were receiving visual feedback during this contraction. During the second contraction, when feedback was not available, torque during both agonist vibration conditions remained comparable to control. In contrast, following the removal of antagonist vibration, which was applied in the first contraction with visual feedback (29.3 %MVC [27.5, 31.1]), torque significantly increased (p = .0026, d = -1.60) compared to control (25.4 %MVC [23.6, 27.2]) during the second contraction. On the contrary, when antagonist vibration was applied in the second contraction without visual feedback (19.9 %MVC [23.2, 26.9]), torque significantly decreased (p < .0001, d = 2.29; see Table 1).

**Table 1.** Torque estimated marginal means, 95% confidence intervals, p-values and Cohen’s d values for each condition at each contraction.

ANTAGONIST VIBRATION ALTERS THE SENSE OF FORCE
| Condition | Mean Torque (%MVC) | 95 % CI | Significance (p-value) | Effect Size (Cohen's d) |
| --- | --- | --- | --- | --- |
| **Contraction 1** |  |  |  |  |
| Control | 23.3 | [21.5, 25.1] | n/a | n/a |
| Agonist vib. w/ C1 | 23.8 | [22.0, 25.6] | .9925 | -0.1852 |
| Agonist vib. w/ C2 | 24.0 | [22.2, 25.8] | .9619 | -0.2867 |
| Antagonist vib. w/ C1 | 23.1 | [21.3, 24.9] | .9997 | 0.0808 |
| Antagonist vib. w/ C2 | 23.1 | [21.3, 24.9] | .9998 | 0.0754 |
| **Contraction 2** |  |  |  |  |
| Control | 25.4 | [23.6, 27.2] | n/a | n/a |
| Agonist vib. w/ C1 | 25.0 | [23.2, 26.8] | .9944 | 0.1714 |
| Agonist vib. w/ C2 | 25.1 | [23.3, 26.9] | .9980 | 0.1324 |
| Antagonist vib. w/ C1 | 29.3 | [27.5, 31.1] | **.0026** | -1.604 |
| Antagonist vib. w/ C2 | 19.9 | [18.1, 21.7] | **< .0001** | 2.295 |

#### Changes from contraction to contraction within each condition

Within the control condition, torque increased (*p* = .04, d = -0.88) from the first contraction (i.e., with visual feedback; 23.3 %MVC [21.5, 25.1]) to the second contraction (i.e., without visual feedback; 25.4 %MVC [23.6, 27.2]). For both agonist vibration conditions, mean torque did not differ significantly across contractions. When antagonist vibration was applied in the first of two contractions with visual feedback, torque significantly increased (*p* < .001, d = -2.57) from the first contraction (23.1 %MVC [21.3, 24.9]) to the second contraction (29.3 %MVC [27.5, 31.1]) following the removal of vibration. In contrast, when antagonist vibration was applied in the second of two contractions without visual feedback, torque significantly decreased (*p* = .002, d = 1.34) from the first contraction (23.1 %MVC [21.3, 24.9]) to the second (19.9 %MVC [18.1, 21.7]; see Table 2 and Figure 3).

**Table 2.** Torque estimated marginal means, 95% confidence intervals, p-values and Cohen’s d values for each contraction within each condition.

| Condition | C1 |  | C2 |  | Significance (p-value) | Effect Size (Cohen's *d*) |
| --- | --- | --- | --- | --- | --- | --- |
|  | Mean Torque (%MVC) | 95 % CI | Mean Torque (%MVC) | 95 % CI |  |  |
| Control | 23.3 | [21.5, 25.1] | 25.4 | [23.6, 27.2] | **.0417** | -0.880 |
| Agonist vib. w/ C1 | 23.8 | [22.0, 25.6] | 25.0 | [23.2, 26.8] | .2229 | -0.523 |
| Agonist vib. w/ C2 | 24.0 | [22.2, 25.8] | 25.1 | [23.3, 26.9] | .2827 | -0.461 |
| Antagonist vib. w/ C1 | 23.1 | [21.3, 24.9] | 29.3 | [27.5, 31.1] | **< .0001** | -2.565 |
| Antagonist vib. w/ C2 | 23.1 | [21.3, 24.9] | 19.9 | [18.1, 21.7] | **.0022** | 1.340 |

### Motor Unit Behaviour

Condition (χ^2^(8) = 113.36, *p* < .0001), contraction (χ^2^(5) = 64.86, *p* < .0001) and their interaction (χ^2^(4) = 63.67, *p* < .0001) were significant predictors of MU firing behaviour.

#### Differences between conditions during each contraction (compared to control)

As expected, MU firing behaviour during the first contraction (i.e., with visual feedback) did not significantly differ between control and any of the vibration conditions. Further, there were no significant differences observed during the second contraction between control and the condition where agonist vibration was applied during contraction 2 without visual feedback. However, when agonist vibration was applied in the first contraction with visual feedback (13.5 pps [12.5, 14.5]), the mean MU firing rate significantly decreased (p = .0003, d = 0.5598) during the second contraction (i.e. following removal of vibration) compared to control (14.5 pps [13.5, 15.5]). Further, following the removal of antagonist vibration applied in the first contraction (i.e., with visual feedback), the mean MU firing rate during the second contraction significantly increased (15.2 pps [14.2, 16.2]; p = .0312, d = -0.3759) compared to control (14.5 pps [12.5, 15.5]), whereas when antagonist vibration was applied in the second contraction (12.8 pps [11.8, 13.8]), the mean MU firing rate was significantly reduced (p < .0001, d = 0.9513; see Table 3).

**Table 3.** Mean MU firing rate estimated marginal means, 95 % confidence intervals, p-values and Cohen’s d values for each condition at each contraction.

| Condition | MU firing rate (pps) | 95 % CI | Significance (p-value) | Effect Size (Cohen's d) |
| --- | --- | --- | --- | --- |
| **Contraction 1** |  |  |  |  |
| Control | 14.2 | [13.2, 15.2] | n/a | n/a |
| Agonist vib. w/ C1 | 14.1 | [13.1, 15.1] | .9925 | 0.0607 |
| Agonist vib. w/ C2 | 13.9 | [13.0, 14.9] | .7720 | 0.1600 |
| Antagonist vib. w/ C1 | 14.1 | [13.1, 15.0] | .9642 | 0.0924 |
| Antagonist vib. w/ C2 | 14.3 | [13.3, 15.2] | .9999 | -0.0193 |
| **Contraction 2** |  |  |  |  |
| Control | 14.5 | [13.5, 15.5] | n/a | n/a |
| Agonist vib. w/ R1 | 13.5 | [12.5, 14.5] | **.0003** | 0.5598 |
| Agonist vib. w/ R2 | 14.1 | [13.1, 15.0] | .3120 | 0.2520 |
| Antagonist vib. w/ R1 | 15.2 | [14.2, 16.2] | **.0312** | -0.3759 |
| Antagonist vib. w/ R2 | 12.8 | [11.8, 13.8] | **< .0001** | 0.9513 |

#### Changes from contraction to contraction within each condition

The mean MU firing rate remained consistent across contractions within the control condition, as well as when agonist vibration was applied in the second contraction. However, when agonist vibration was applied during the first contraction (i.e., with visual feedback), the mean MU firing rate significantly decreased (*p* = .012, d = 0.3433) from contraction one (14.1 pps [13.1, 15.2]) to two (13.5 pps [12.5, 14.5]). Further, antagonist vibration applied in the first of two contractions caused a significant increase (*p* < .0001, d = -0.6241) in the mean MU firing rate from the first (14.1 pps [13.1, 15.0]) to the second contraction (15.2 pps [14.2, 16.2]). On the other hand, antagonist vibration applied in the second of two contractions caused a significant decrease (*p* < .0001, d = 0.8148) in the mean MU firing rate from the first (14.3 pps [13.3, 15.2]) to second contraction (12.8 pps [11.8, 13.8]; see Table 4 and Figure 4).

**Table 4.** Mean MU firing rate estimated marginal means, 95 % confidence intervals, p-values and Cohen’s d values for each contraction within each condition.

| Condition | C1 | | C2 | | Significance<br>($p$-value) | Effect Size<br>(Cohen's<br>$d$) |
| --- | --- | --- | --- | --- | --- | --- |
|  | Mean<br>MU<br>Firing<br>(pps) | 95 % CI | Mean MU<br>Firing<br>(pps) | 95 % CI |  |  |
| Control | 14.2 | [13.2, 15.2] | 14.5 | [13.5, 15.5] | .2428 | -0.1558 |
| Agonist vib. w/ C1 | 14.1 | [13.1, 15.2] | 13.5 | [12.5, 14.5] | **.0125** | 0.3433 |
| Agonist vib. w/ C2 | 13.9 | [13.0, 14.9] | 14.1 | [13.1, 15.0] | .6315 | -0.0638 |
| Antagonist vib. w/ C1 | 14.1 | [13.1, 15.0] | 15.2 | [14.2, 16.2] | **< .0001** | -0.6241 |
| Antagonist vib. w/ C2 | 14.3 | [13.3, 15.2] | 12.8 | [11.8, 13.8] | **< .0001** | 0.8148 |

**Figure 4.**
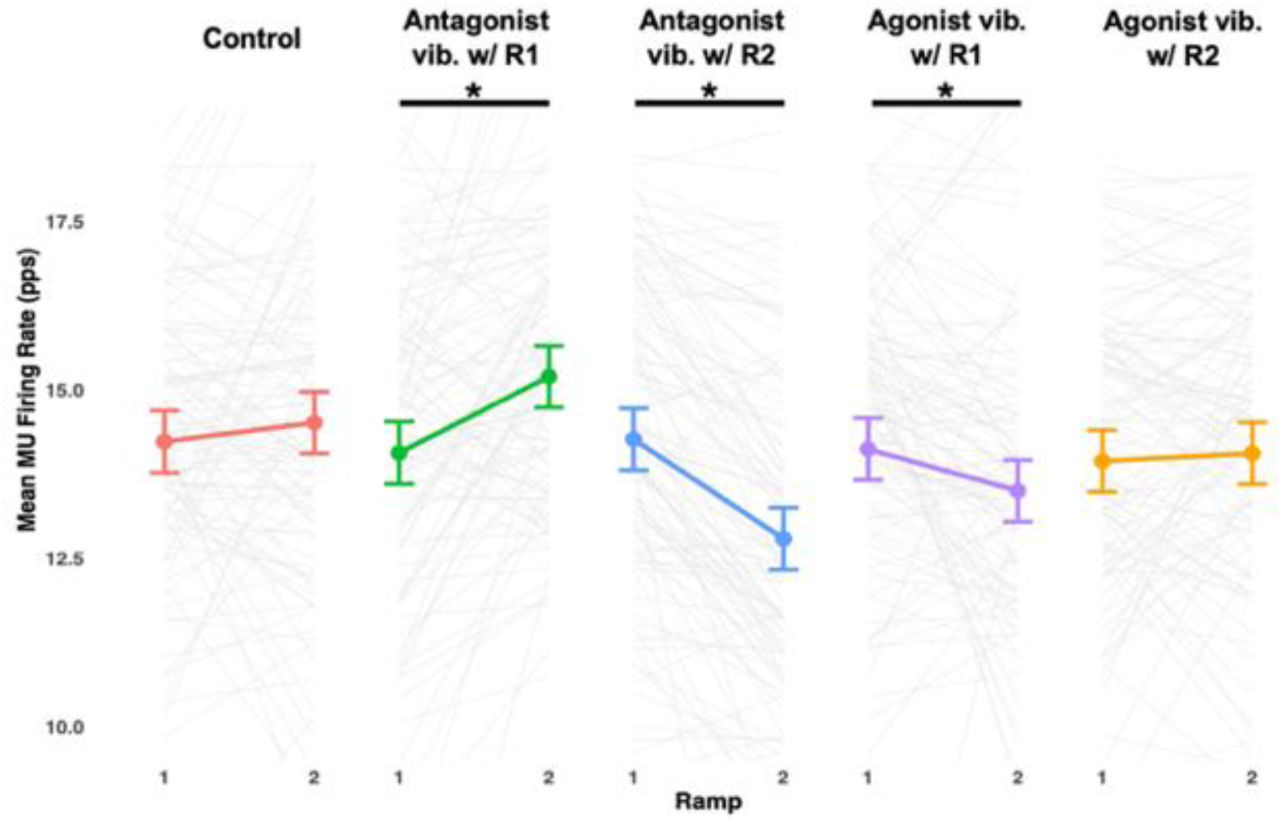
Mean motor unit firing rate (pps) values for the tibialis anterior (TA) during each contraction within each experimental condition. Points represent the model estimates from the linear mixed effects model and the error bars are the corresponding 95% confidence intervals. Grey lines in the background of each condition correspond to changes in firing behaviour for each unit identified. Asterisks indicate significant difference in mean torque between contractions (p < .05).

#### Torque reproduction errors *Absolute error*

Contraction (χ^2^(5) = 17.47, *p* = .004) was a significant predictor of AE, although condition (χ^2^(8) = 14.85, *p* = .062) and the interaction between contraction and condition (χ^2^(4) = 5.06, *p* = .282) were not. No significant differences in AE were observed between control and any vibration condition during either contraction (see Table 5). Across conditions, AE remained similar from the first to the second contraction for the control and both agonist vibration conditions. In contrast, both antagonist vibration conditions resulted in a significant increase in AE across contractions. When applied during the first contraction, AE increased (*p* = .009, d = -1.122) from the first (9.28 % [3.80, 14.8]) to the second contraction (18.61 % [13.1, 24.1]) and when applied during the second contraction, AE increased (*p* = .002, d = -1.338) from the first (9.15 % [3.67, 14.6]) to the second (20.28 % [14.80, 25.8]) contraction (see Table 6 and Figure 5).when antagonist vibration was applied in the first contraction, AE significantly increased (p =, d =) from the first (9.28 % [3.80, 14.8]) to the second contraction (18.61 % [13.1, 24.1]). Further, when antagonist vibration was applied in the second contraction, AE increased (d =) from the see figure 5) for both antagonist vibration conditions

**Table 5.** AE estimated marginal means, 95 % confidence intervals, p-values and Cohen’s d values for each condition during each contraction.

| Condition | AE (%) | 95 % CI | Significance (p-value) | Effect Size (Cohen's d) |
| --- | --- | --- | --- | --- |
| **Contraction 1** |  |  |  |  |
| Control | 9.02 | [3.54, 14.5] | n/a | n/a |
| Agonist vib. w/ C1 | 7.52 | [2.04, 13.0] | .9933 | 0.1800 |
| Agonist vib. w/ C2 | 7.04 | [1.56, 12.5] | .9810 | 0.2368 |
| Antagonist vib. w/ C1 | 9.28 | [3.80, 14.8] | 1.000 | -0.0314 |
| Antagonist vib. w/ C2 | 9.15 | [3.67, 14.7] | 1.000 | -0.0160 |
| **Contraction 2** |  |  |  |  |
| Control | 13.14 | [7.66, 18.6] | n/a | n/a |
| Agonist vib. w/ C1 | 10.06 | [4.58, 15.5] | .9071 | 0.3711 |
| Agonist vib. w/ C2 | 9.52 | [4.04, 15.0] | .8457 | 0.4349 |
| Antagonist vib. w/ C1 | 18.61 | [13.1, 24.1] | .5388 | -0.6569 |
| Antagonist vib. w/ C2 | 20.28 | [14.8, 25.8] | .2680 | -0.8579 |

**Table 6.**
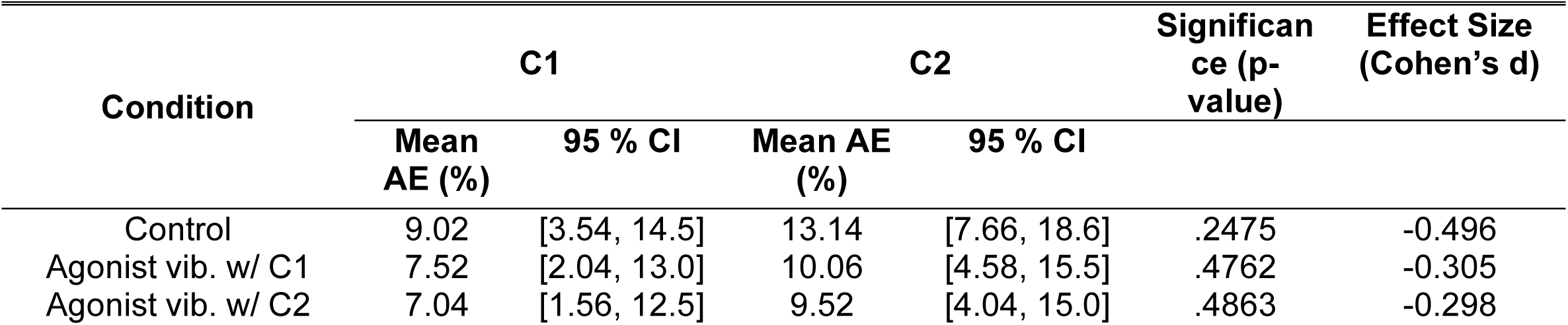
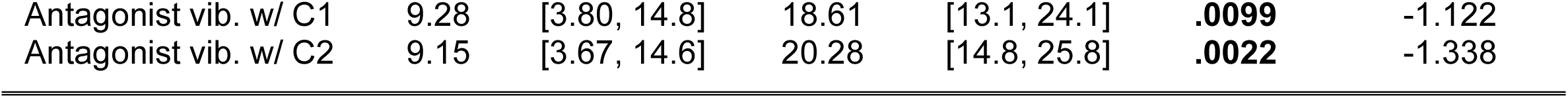
AE estimated marginal means, 95 % confidence intervals, p-values and Cohen’s d values for each contraction within each condition.

| Condition | C1 |  | C2 |  | Significance (p-value) | Effect Size (Cohen's d) |
| --- | --- | --- | --- | --- | --- | --- |
|  | Mean AE (%) | 95 % CI | Mean AE (%) | 95 % CI |  |  |
| Control | 9.02 | [3.54, 14.5] | 13.14 | [7.66, 18.6] | .2475 | -0.496 |
| Agonist vib. w/ C1 | 7.52 | [2.04, 13.0] | 10.06 | [4.58, 15.5] | .4762 | -0.305 |
| Agonist vib. w/ C2 | 7.04 | [1.56, 12.5] | 9.52 | [4.04, 15.0] | .4863 | -0.298 |
| Antagonist vib. w/ C1 | 9.28 | [3.80, 14.8] | 18.61 | [13.1, 24.1] | **.0099** | -1.122 |
| Antagonist vib. w/ C2 | 9.15 | [3.67, 14.6] | 20.28 | [14.8, 25.8] | **.0022** | -1.338 |

**Figure 5.**
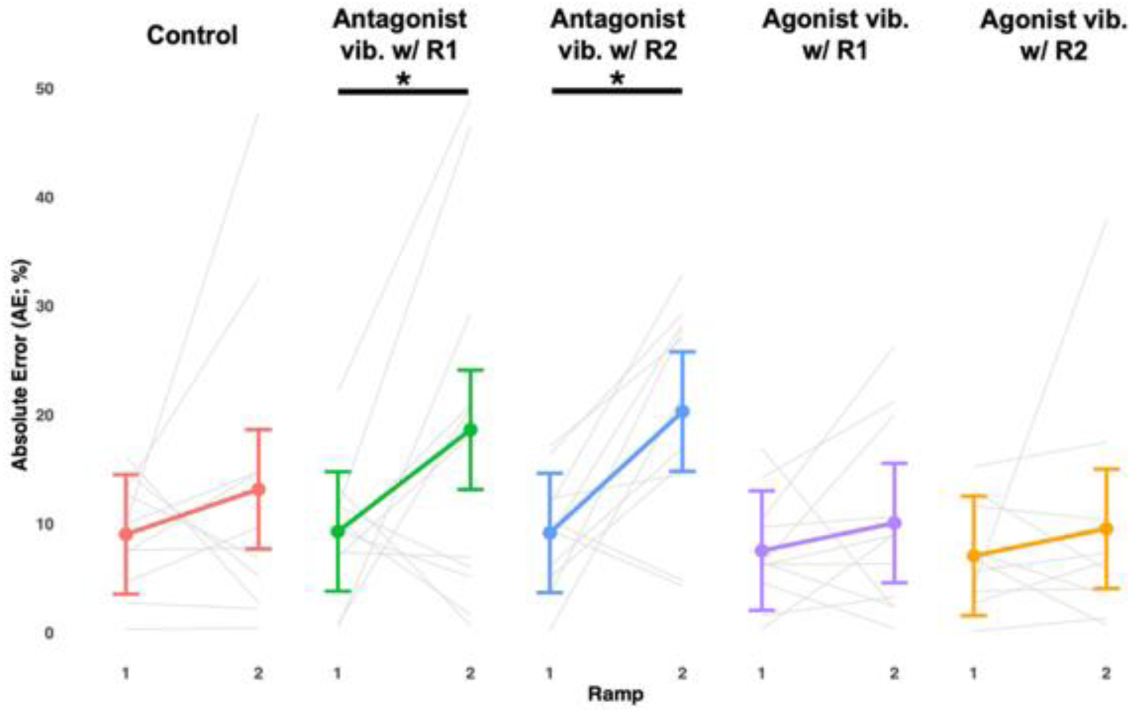
Mean absolute torque reproduction error (AE; %) values during each contraction within each experimental condition. Points represent the model estimates from the linear mixed effects model and the error bars are the corresponding 95% confidence intervals. Grey lines in the background of each condition correspond to individual participant AE. Asterisks indicate significant difference in mean torque between contractions (p < .05).

#### Constant error

Contraction (χ^2^(5) = 42.43, *p* < .0001), condition (χ^2^(8) = 61.78, *p* < .0001) and their interaction (χ^2^(4) = 35.35, *p* < .0001) were significant predictors of CE. During the first contraction, CE remained similar to control for all vibration conditions and all values were slightly negative, indicating a consistent undershoot. During the second contraction, there were no major differences in CE between control and either of the agonist vibration conditions (see Table 7). Further, there were no major differences between contractions within either agonist vibration condition (see Table 8). In contrast, CE differed significantly across contractions within the control and two antagonist vibration conditions. Within the control condition, CE increased (*p* = .0417, d = - 0.880) from an undershoot during the first contraction (-6.71 % [-13.9, 0.476]) to a slight overshoot during the second contraction (1.72 % [-5.46, 8.91]). When antagonist vibration was applied during the first contraction, CE significantly increased (*p* < .0001, d = -2.57) from a slight undershoot in the first contraction (-7.49 % [-14.7, -0.299]) to a drastic overshoot during the second contraction (17.11 % [9.93, 24.3]). Conversely, when antagonist vibration was applied during the second contraction only, CE significantly decreased (*p* = .0022, d = -2.22) from a slight undershoot in the first contraction (-7.43 % [-14.6, -0.247]) to a severe undershoot in the second contraction (-20.28 % [-27.5, -13.1]; see Figure 6).

**Table 7.** CE estimated marginal means, 95 % confidence intervals, p-values and Cohen’s d values for each condition during each contraction.

| Condition | CE (%) | 95 % CI | Significance (p-value) | Effect Size (Cohen's d) |
| --- | --- | --- | --- | --- |
| **Contraction 1** |  |  |  |  |
| Control | -6.71 | [-13.9, 0.476] | n/a | n/a |
| Agonist vib. w/ C1 | -4.94 | [-12.1, 2.25] | .9925 | -0.1852 |
| Agonist vib. w/ C2 | -3.96 | [-11.2, 3.23] | .9619 | -0.2867 |
| Antagonist vib. w/ C1 | -7.49 | [-14.7, -0.299] | .9997 | 0.0808 |
| Antagonist vib. w/ C2 | -7.43 | [-14.6, -0.247] | .9998 | 0.0754 |
| **Contraction 2** |  |  |  |  |
| Control | 1.72 | [-5.46, 8.91] | n/a | n/a |
| Agonist vib. w/ C1 | 0.08 | [-7.11, 7.27] | .9944 | 0.1714 |
| Agonist vib. w/ C2 | 0.455 | [-6.73, 7.64] | .9980 | 0.1324 |
| Antagonist vib. w/ C1 | 17.11 | [9.93, 24.3] | **.0026** | -1.605 |
| Antagonist vib. w/ C2 | -20.28 | [-27.5, -13.1] | **< .0001** | -2.295 |

**Table 8.**
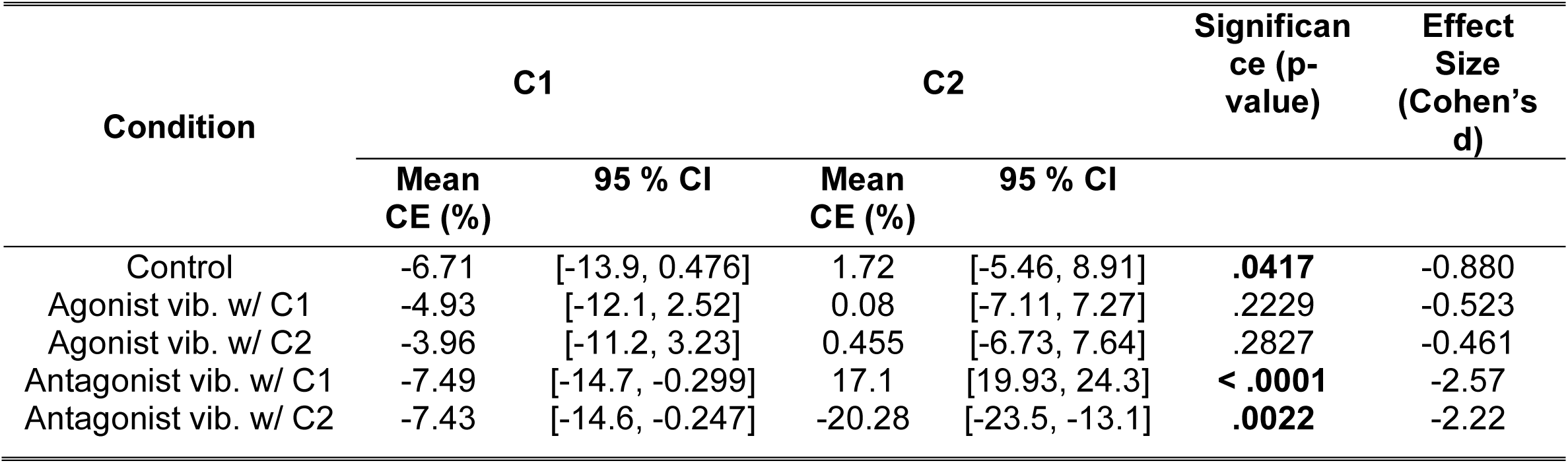
CE estimated marginal means, 95 % confidence intervals, p-values and Cohen’s d values for each contraction within each condition.

**Figure 6.**
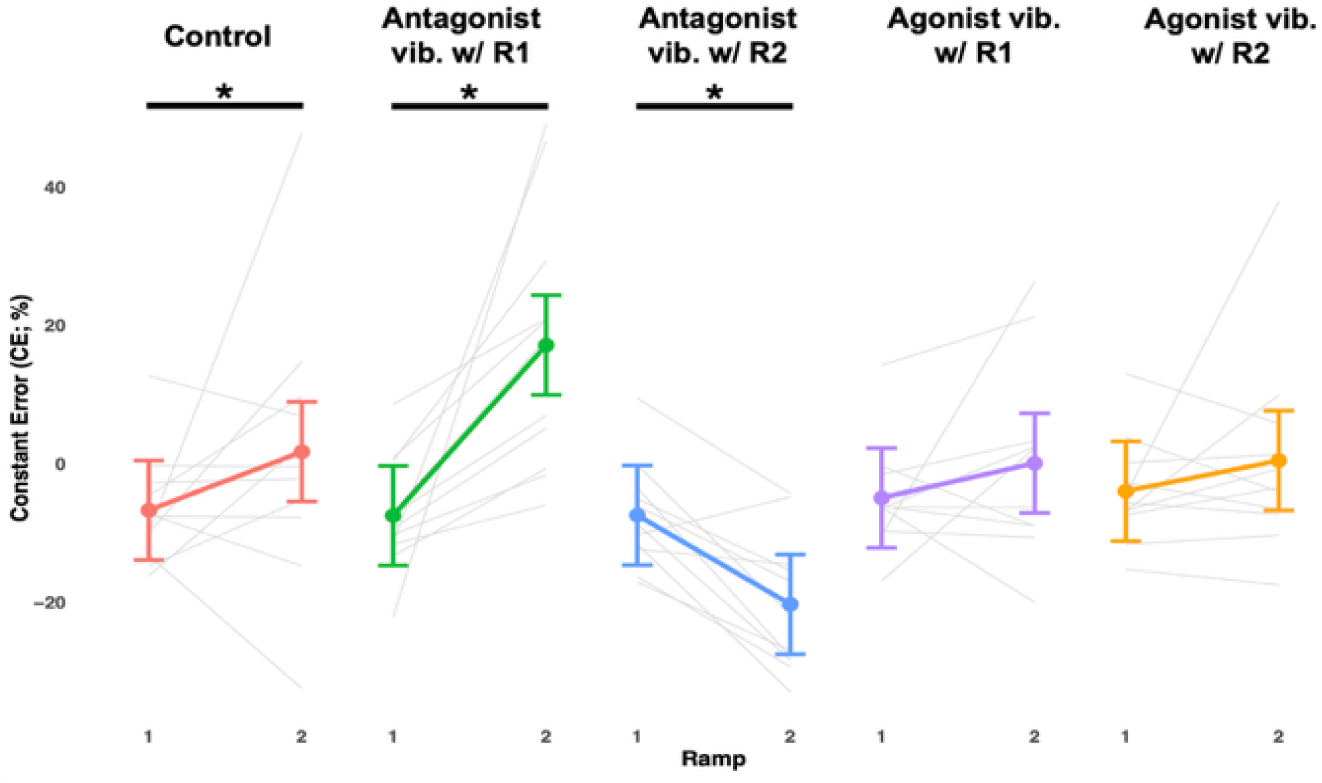
Mean constant torque reproduction error (CE; %) values during each contraction within each experimental condition. Points represent the model estimates from the linear mixed effects model and the error bars are the corresponding 95% confidence intervals. Grey lines in the background of each condition correspond to individual participant CE. Asterisks indicate significant difference in mean torque between contractions (p < .05).

## DISCUSSION

The present study aimed to quantify the effects of both antagonist and agonist tendon vibration on the ability to reproduce torque under conditions of matched effort (i.e., neural drive). In doing so, we indirectly assessed the effects of sensory feedback designed to alter motoneuronal gain on torque output in conditions where voluntary effort was held constant. The application and removal of antagonist tendon vibration elicited much larger effects on the ability to reproduce torque, error and MU firing properties than did agonist tendon vibration. Our findings demonstrate a consistent undershoot relative to target torque (i.e., under production of torque) during contractions where antagonist tendon vibration was applied and an overshoot relative to target torque (i.e., over production of torque) during contractions immediately following the removal of antagonist tendon vibration. These deviations were accompanied by large AE values, as well as large directional shifts in CE values (i.e. negative shift during applied antagonist vibration; positive shift post-antagonist vibration), reinforcing reduced torque reproduction accuracy. On the other hand, we observed small and inconsistent effects on torque generation during both contractions with applied agonist vibration and contractions immediately following the removal of agonist tendon vibration. Consistent with this finding, agonist tendon vibration produced negligible changes in both AE and CE values, suggesting little impact on torque reproduction accuracy. These findings support our hypothesis that application of antagonist tendon vibration would transiently alter the ability to reproduce target torque, whereas vibration applied to the agonist tendon would have a reduced impact on torque reproduction. Importantly, the behavioural differences observed across experimental conditions were reflected in the underlying MU firing patterns, offering insight into neural mechanisms which may contribute to these effects.

The substantial effects on torque reproduction observed with manipulation of antagonist tendon vibration and lack thereof with manipulation of agonist tendon vibration are supported by the corresponding underlying MU behaviour. Specifically, similar to torque, mean MU firing rate was markedly reduced during contractions with applied antagonist tendon vibration and increased during contractions immediately after the removal of antagonist vibration. In contrast, agonist tendon vibration did not substantially alter MU firing during its application. However, somewhat unexpectedly, mean MU firing rate significantly decreased during the contraction following the removal of agonist tendon vibration, highlighting that distinct neural mechanisms likely underlie the competing effects of antagonist versus agonist tendon vibration.

### Potential mechanisms differentiating antagonist versus agonist vibration

#### Reciprocal inhibition reduces magnitudes of PICs

The pronounced alterations in torque, error and MU firing observed with manipulation of antagonist, but not agonist tendon vibration suggest that reciprocal inhibition is likely a key mechanism at play. Reciprocal Ia inhibition is a spinal mechanism in which Ia afferents from the agonist muscle activate Ia inhibitory interneurons, which in turn, inhibit the antagonist motoneuron pool (Baldissera et al., 1981; Pierrot-Deseilligny & Burke, 2005). When high-frequency vibration is applied to the antagonist muscle tendon, activation of muscle spindles increases Ia afferent discharge, which causes inhibitory postsynaptic potentials (IPSPs) onto agonist motoneurons, effectively reducing their net excitatory drive (i.e. gain; Burke et al., 1976). Reciprocal Ia inhibition has also been shown to markedly suppress magnitudes of PICs, which when activated, play an integral role in amplifying synaptic input and sustaining motoneuron firing (Heckman et al., 2008). In nonhuman animal models, small increases in reciprocal Ia inhibition elicited by activation of muscle spindles via antagonist muscle stretch reduced PIC estimates approximately 2-fold (Hyngstrom et al., 2007). Likewise, electrical stimulation-induced measures of reciprocal Ia inhibition have been suggested to correlate with the magnitude of PICs estimated from motor units in humans (Vandenbark & Kalmar, 2014). Emerging evidence further supports the modulatory effects of reciprocal Ia inhibition on estimates of PICs in humans. For instance, Mesquita et al., (2022) demonstrated decreased estimates of PIC amplitude in the gastrocnemius with low-frequency stimulation of the common peroneal nerve, and similar reductions in estimates of PICs were observed in our previous work examining both the plantarflexors and dorsiflexors with high-frequency vibration applied to their respective antagonist muscle tendons (Pearcey et al., 2022). These findings collectively suggest that increased Ia input from the antagonist suppresses PIC expression and therefore the intrinsic excitability of motoneurons and provide a mechanistic explanation for motor behaviours exhibited throughout the present study.

When participants completed the force reproduction task, applied antagonist tendon vibration resulted in a severe torque undershoot relative to the target contraction as well as reduced corresponding MU firing rates. On the other hand, when participants attempted to reproduce torque generated during the contraction with applied vibration following the removal of that vibration, they severely overshot compared to target and MU firing largely increased. This aligns with established mechanisms discussed above. Since PICs amplify synaptic input and sustain motoneuronal firing for a given level of descending drive, their suppression due to vibration would be expected to reduce motoneuronal gain, which would in turn, reduce torque output despite constant effort, which is precisely what occurred in the present study. Removal of this vibration would be expected to evoke a rebound in motoneuronal gain, since reciprocal inhibition is withdrawn with the removal of vibration, meaning PICs would once again be expressed as normal. Because participants were instructed to match percieved effort that they generated within the applied vibration contraction, re-expression of normal PIC magnitudes would be expected to produce greater torque and MU firing for the same voluntary drive, which again, is what occurred within the present study. This consistent pattern of torque undershoot during antagonist vibration and torque overshoot following its removal, and the aligning MU firing behaviour, supports that transient modulation of PICs via reciprocal inhibition likely played a key role in the impairments in force reproducibility and underlying MU behaviour observed throughout the present study.

#### Ia input from the agonist does not affect magnitudes of PICs

In contrast to the large inhibitory influence of Ia input from the antagonist muscle on the agonist motor pool, increased Ia excitation from the agonist muscle seemed to have little meaningful impact on torque, error and MU firing. Opposite to reciprocal Ia inhibition, Ia afferents from the agonist provide direct excitatory monosynaptic input to the alpha motoneurons of the same muscle, enhancing excitatory drive to the agonist motoneuron pool (Pierrot-Deseilligny & Burke, 2005). In reduced animal preparations, Ia excitatory input is sufficient to initiate PIC activation and trigger self-sustained firing of motoneurons, but once PICs become activated (i.e. under conditions of ongoing voluntary contraction, for instance), additional Ia input has a negligible effect on firing behaviour (Heckman & Lee, 1999). This aligns with findings of the present study, which revealed only modest and inconsistent effects of agonist vibration on force reproduction, error, and MU behaviour regardless of how it was manipulated within the task.

Despite the substantial increase in homonymous Ia afferent discharge and the additional reflexive excitation potentially introduced by the tonic vibration reflex with agonist vibration, neither torque nor MU firing changed meaningfully. This was the case even though we specifically instructed participants to ensure that their effort (i.e., voluntary drive) remained the same in the presence and absence of vibration, suggesting that PICs, or gain control, remained constant throughout periods of agonist vibration. If Ia afferent input from the agonist suppressed PICs during voluntary contraction, a reduction in torque and MU firing would have been expected, similar to the clear decrease observed when the antagonist was vibrated. On the other hand, if Ia excitation from the agonist had amplified PICs and therefore motoneuronal gain, an increase in torque and MU firing would have been expected. Since torque and MU firing remained relatively stable (i.e. no consistent and significant increases or decreases), it seems reasonable to assume that PIC amplitude remained unchanged. Although this does not rule out the possibility that descending drive and monoaminergic facilitation may be slightly adjusted as a result of agonist vibration, as proposed by Lapole et al., (2023), these types of adjustments did not impact torque generation nor MU firing in the present study due to the nature of the task.

### Methodological Considerations

There are several methodological limitations which should be addressed. First, we only tested at one submaximal intensity, which was 25% of each participant’s maximum, and from only one muscle, which was the TA (i.e. the primary ankle dorsiflexor). Therefore, we cannot confirm whether effects induced by antagonist tendon vibration are generalizable to force levels that are below or exceed this intensity, or in other muscles. Given that our chosen intensity is relatively low, it is likely that smaller, lower-threshold motoneurons are primarily recruited rather than larger and higher-threshold motoneurons (Henneman, 1957). Since smaller motoneurons have higher input resistance than their larger counterparts (Henneman, Somjen & Carpenter, 1965), it is possible that they are subject to a greater relative influence of PICs. Though this idea has not been extensively investigated in humans, findings from animal models suggest that smaller motoneurons may exhibit stronger intrinsic amplification than larger motoneurons and consequently possess proportionally larger contributions of PICs relative to cell size (Huh et al., 2017). Therefore, vibration-related PIC modulation may be particularly evident at lower force levels.

Second, we did not assess estimates of PICs directly in this study, meaning any conclusions made surrounding their contributions are indirect. However, our experimental design and interpretation are grounded in well-established physiological principles indicating that PICs are highly sensitive to Ia afferent input from the antagonist. Further, our findings closely align with established PIC modulation patterns during antagonist tendon vibration in both nonhuman animals (Hyngstrom et al., 2007) as well as humans (Mesquita et al., 2022; Pearcey et al., 2022). Therefore, while direct measurements of PIC magnitudes would have been beneficial, the indirect evidence provided here remains robust and consistent with existing work.

Next, utilization of HD-sEMG allows for sampling of only a portion of the motor pool since it is inherently biased towards MUs with larger muscle fibres and those that are closer to the surface of the skin (Farina et al., 2014). It is therefore possible that additional units which were not identified could have contributed to the effects of antagonist tendon vibration. However, compared to single channel or bipolar EMG setups, decomposition of HD-sEMG signals yields a much larger concurrent sample of MUs, providing a more robust window into overall motor behaviour (Farina and Holobar, 2016). Even though we collected only a sample of active MUs, this subset is widely considered representative of the behaviour of the entire motor pool since MUs receive common synaptic input (Hug et al., 2023). As a result, changes in firing properties of identified units can be interpreted as proportional changes in the common synaptic input to the whole motor pool (Del Vecchio et al., 2020). This represents the most reliable, but also non-invasive, insight into motoneuronal behaviours currently available.

Finally, we are uncertain of the effects of agonist and antagonist vibration on perceptual centers in the brain. In the current study, participants were instructed to maintain consistent effort from the first to second contraction in each of the force reproduction tasks. However, it remains possible that vibration could have influenced perceptual or cortical processes, meaning that some of the observed effects of vibration could have potentially originated supraspinally rather than exclusively at the spinal level. Further investigation implementing electroencephalography or neurostimulation paradigms would help clarify this distinction.

## CONCLUSION

Our findings demonstrate that antagonist, but not agonist, tendon vibration severely disrupts the ability to reproduce force, and that these behavioural impairments closely align with underlying MU firing patterns. During the force reproduction task, these impairments manifested as reductions in torque when antagonist vibration was applied in the second of two contractions without visual feedback and increases in torque following the removal of antagonist vibration applied in the first of two contractions with visual feedback. Agonist vibration on the other hand, had minimal and inconsistent effects on both force reproducibility and error, regardless of how it was manipulated within the task. Both the magnitude and the direction of the observed errors in force reproduction in response to antagonist vibration align with the pre-established impact of reciprocal Ia inhibition on PICs. In contrast, additional Ia excitation from the agonist produced no meaningful and consistent changes in torque reproduction behaviour nor MU firing, which reinforces the insensitivity of PICs to additional excitatory Ia input once they are already activated. The results of the present study therefore strongly support reciprocal Ia inhibition via antagonist tendon vibration as a means of altering force reproduction capabilities, since it severely diminishes the intrinsic gain of motoneurons. This highlights PIC modulation as a central mechanism which contributes to shaping the sense of force under altered proprioceptive conditions.

## Supplementary Table

### Effort ratings during each contraction of each force reproduction pair

**Table 9.** Force reproduction task percieved effort ratings. C1 = contraction one within each force reproduction pair and C2 = contraction 2 within each force reproduction pair.

| Participant | CONTROL |  | VIB W/ C1 |  |  |  | VIB W/ C2 |  |  |  |
| --- | --- | --- | --- | --- | --- | --- | --- | --- | --- | --- |
|  | No Vibration |  | Antagonist |  | Agonist |  | Antagonist |  | Agonist |  |
|  | C1 | C2 | C1 | C2 | C1 | C2 | C1 | C1 | C1 | C2 |
| **M01** | 2 | 2 | 3 | 3 | 2 | 2 | 2 | 2 | 2 | 2 |
| **M02** | 3 | 3 | 4 | 4 | 4 | 4 | 4 | 4 | 3 | 3 |
| **M03** | 4.5 | 4.5 | 4 | 4 | 4.5 | 4.5 | 4 | 4 | 4 | 4 |
| **M04** | 2 | 2 | 3 | 3 | 3 | 3 | 2 | 2 | 2 | 2 |
| **M05** | 2 | 2 | 2 | 2 | 2 | 2 | 2 | 2 | 2 | 2 |
| **M06** | 3 | 3 | 4 | 4 | 4 | 4 | 3 | 3 | 3 | 3 |
| **F01** | 2 | 2 | 2 | 2 | 2 | 2 | 2 | 2 | 2 | 2 |
| **F02** | 5 | 5 | 5 | 5 | 5 | 5 | 6 | 6 | 5 | 5 |
| **F03** | 3 | 3 | 4 | 4 | 3 | 3 | 3 | 3 | 4 | 4 |
| **F04** | 4 | 4 | 5 | 5 | 4 | 4 | 4 | 4 | 3 | 3 |
| **F05** | 1 | 1 | 3 | 3 | 2 | 2 | 3 | 3 | 4 | 4 |

